# Freshwater fish egg dispersal by attaching to waterbirds

**DOI:** 10.1101/2024.03.11.584339

**Authors:** Akifumi Yao, Miyuki Mashiko, Yukihiko Toquenaga

## Abstract

Dispersal over geographic barriers plays an essential role in colonization, gene flow, metapopulation dynamics, and invasion (Bowler and Benton 2005). Since dry lands strictly separate freshwater habitats, as expressed by the phrase “islands of water in a sea of dry land” (Faulks, Gilligan, and Beheregaray 2010), dispersal among freshwater waterbodies by themselves is challenging of aquatic organisms. Rumors have existed worldwide that freshwater fish eggs are dispersed by attaching to (ectozoochory) or excretion from (endzoochory) waterbirds (Hirsch et al. 2018). It is well documented that waterbirds disperse aquatic plants, zooplankton, and various aquatic invertebrates, which are co-distributed with fishes (Green et al. 2023). However, there is only two reported cases of empirical evidence of endzoochory in freshwater fishes and no scientific evidence of ectozoochory (Hirsch et al. 2018; Silva et al. 2019; Lovas-Kiss et al. 2020; Green et al. 2023). Here, we show that the southern medaka (*Oryzias latipes*, hereafter medaka) egg can travel passively by attaching to waterbirds.

## INTRODUCTION

Medaka is a small freshwater fish living in the shallows of ponds, lakes, paddy fields, and rivers in Japan except Hokkaido Island (Senou 2013). It lays eggs on various substrates, such as submerged aquatic plants (Iwata et al. 2015). The picture book “Flying Medaka in the Sky (“*Soratobu medaka*” in Japanese) ” reported that shoals of medaka were found in a shallow concrete pool that was used to wash trucks. The writer proposed that waterbirds such as night herons (*Nycticorax nycticorax*) carried medaka eggs from a nearby stream because he thought that only birds could reach both the concrete pool and the creek, and he was convinced that such waterbirds carried eggs attached to aquatic plants tangled on their feet (Nakamura 1999). However, this scenario has so far not been validated scientifically.

## RESULTS

To test this scenario, we first conducted a field experiment to examine whether waterbirds carry submerged aquatic plants that serve as medaka’s spawning substrates. We constructed two experimental ponds about 1 m × 2 m among rice paddy fields (Hojo, Tsukuba, Ibaraki, Japan; Fig. 1a; Appendix S1) to which herons and egrets fly for foraging throughout the year. To prevent genetic disturbance of local populations, we used artificial aquatic plants to substitute for submerged aquatic plants and did not attach medaka’s egg. Thirty-six artificial plants were placed in one of the ponds (source pond, right one in Fig. 1a) and we observed whether the waterbirds dispersed them to the other pond (sink pond, left one in Fig. 1a) by motion capture camera traps.

**Figure 1.**
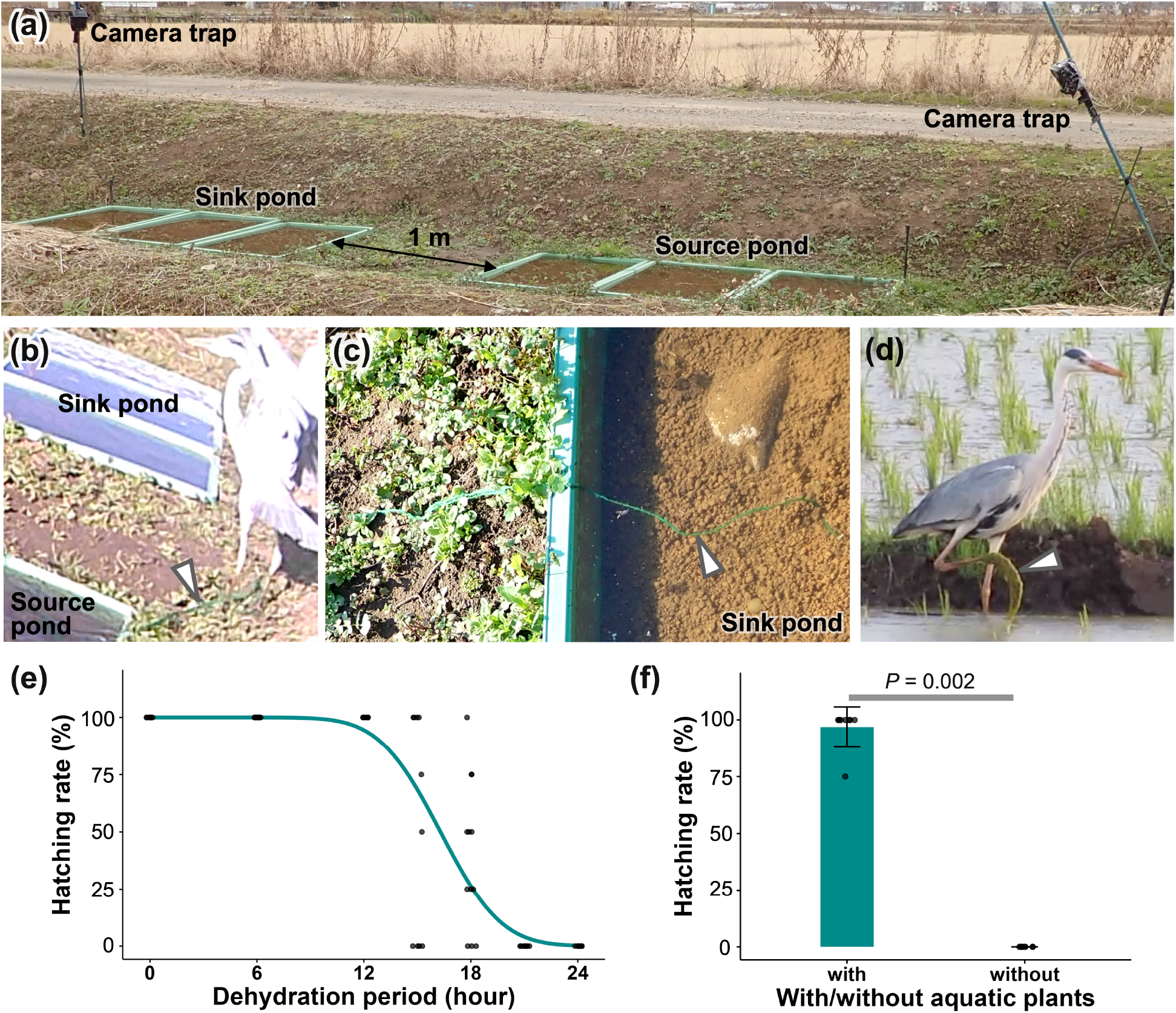
Field and laboratory experiments showed that medaka eggs could be dispersed passively by attaching to waterbirds. (a) Field experimental system (b) A grey heron hooked artificial aquatic plants on its foot and flew away. Arrowhead shows artificial aquatic plants (clipped from video S2). (c) An artificial aquatic plant was found at the sink pond. Arrowhead shows the artificial aquatic plant (December 14, 2019). (d) A grey heron carried clumps of algae on its foot and walked in paddy fields. Arrowhead shows algae on heron foot (clipped from video S3). (e) Hatching rates of eggs with attaching aquatic plants (N = 7–12 biological replicates with four eggs in each replicate). (f) Aquatic plants enhance survivability of eggs in the air (N = 8 biological replicates with four eggs in each replicate). Photo credit: (a-c) Akifumi Yao, (d) Miyuki Mashiko.

On December 5, 2019, the camera trap recorded that a grey heron (*Ardea cinere*a) hooked the artificial plants on its foot and flew away from the sink pond (Fig. 1b, Video S1, 2). The artificial aquatic plants were observed around experimental ponds that were located up to 6 m from the source pond (observed on December 8). In addition, an artificial aquatic plant was found in the sink pond on another day (Fig. 1c, observed on December 14). Although the detailed traveling process of the plant was not recorded, one heron walked from source pond to sink pond before this observation. These results suggests that the artificial aquatic plants were carried by such herons. Furthermore, we happened to observe a grey heron walking between paddy fields with a clump of algae on it foot near experimental ponds in Hojo (Fig. 1d, Video S3, recorded May 23, 2019). These observations suggest aquatic plants can be dispersed passively by attaching to waterbirds, and medaka eggs can be carried between waterbodies if they attach to such aquatic plants.

Next, we examined whether medaka eggs attached to aquatic plants could survive in the air during dispersal by birds. Clutches of eggs were attached to 5-cm pieces of aquatic plants (*Egeria densa*). Then, they were exposed to the air for 0–24 hours. After the treatment, eggs were reared, and hatching rates were calculated (Appendix S1). When eggs were attached to aquatic plants, they could survive in the air for up to 18 hours and then hatch, with a median lethal period of 16.3 ± 0.3 hours (probit regression, Fig. 1e). In contrast, all eggs died when they were in the air for 6 hours without being attached to aquatic plants (Fig. 1f, hatching rate of eggs with aquatic plants: 96.9%, that of without aquatic plants: 0%, Wilcoxon signed-rank test, *Z*=3.051, *P*=0.002). These results suggest that, although medaka eggs do not have strong desiccation tolerance, their eggs could survive out of the water for a certain period by attaching to moist aquatic plants.

## DISCUSSION

Our experiments suggest that medaka eggs attached to aquatic plants can be dispersed by waterbirds and the scenario proposed by the picture book may be true. Previous studies reported that seeds of various aquatic plants and plant bodies of small floating aquatic plants could be dispersed by bird-mediated ectozoochory (Coughlan et al. 2017; Coughlan, Kelly, and Jansen 2017). However, there have been no reports of ectozoochory on plant bodies of submerged aquatic plants that serve as spawning substrates for medaka. Our observations suggest that plant bodies of submerged macrophytes can also be dispersed by ectozoochory, although it has been reported that waterbirds carry seeds of such aquatic plants among continents (Viana et al. 2013). In this study, the dispersal of aquatic plants was observed in December, which is not the reproductive season of medaka. However, herons and egrets fly to shallow waterbodies, which are suitable habitats for medaka, for foraging throughout the year, including medaka’s reproductive season from April to September (Iwamatsu 2004). Therefore, dispersal of aquatic plants can occur during the reproductive season of medaka.

This study, together with previous studies, suggests that waterbirds can carry eggs of various freshwater fishes. Previous research showed that killifish eggs dispersed by coscroba swans (*Coscoroba coscoroba*) (Silva et al. 2019). Killifish spawn eggs at the bottom of temporal ponds in deserts, and their eggs have a strong chorion. Furthermore, two carp species’ (*Carpio carpio and C. gibelio*) eggs with soft chorion are spawned on underwater substrates can survive and be dispersed through the birds’ gut (Lovas-Kiss et al. 2020). Our study demonstrated that medaka eggs spawned on aquatic plants can be dispersed by attaching to waterbirds. Also, based on surveys of artificial ponds, Garcia et al. 2023 suggested that waterbirds may disperse European perch’s eggs (*Perca fluviatilis*). Taken together, these findings show that various freshwater fishes spawned in shallow water can be dispersed both by endzoochory and ectozoochory, suggesting that birds potentially disperse more various other freshwater fishes.

Although it is expected that various freshwater fishes could disperse by waterbirds, unique reproduction features of medaka can accelerate frequencies of passive dispersal by waterbirds. First, medaka spawned eggs on substrates. They spawned on various substrates such as submerged aquatic plants, floating aquatic plants, emergent plants, bryophytes, algae, bricks of wood, and fallen leaves (Kobayashi et al. 2012; Iwata et al. 2015). It is thought that eggs attached to the substrate are more frequently trapped on the waterbird’s body than eggs alone. In addition, wet substrates keep eggs moist and improve survivability during dispersal (Fig. 1e, f), suggesting that this feature could increase the ease of dispersal by waterbirds. Second, field observations and laboratory work have shown that this fish prefers laying their eggs on substrates near the water surface rather than near the bottom (Kobayashi et al. 2012; Kamide, Kominami, and Kobayashi 2017). Although this feature is thought to be adaptive to avoid predation or oxygen deficiency (Kobayashi et al. 2012), it may also increase the possibility of contact with birds.

Overall, this study provides the first case of the ectozoochory of freshwater fish eggs by waterbirds and illustrates how animals can cross barriers and disperse using other animals as vectors. Also, we hypothesize that the frequencies of waterbird-mediated dispersal affect freshwater fish populations. Freshwater fishes spawn their eggs in various seasons and on various substrates. Such variations are thought to affect the frequencies of bird-mediated dispersal and the differences of these frequencies among species. If so, it is expected that differences in reproductive traits may lead to differences in the ease of dispersal by waterbirds and, in the long term, to differences in the population dynamics among freshwater fishes. We hope future population genetic studies will help to elucidate how waterbirds contribute to assemblies of fish populations in freshwater habitats.

## ACKNOWLEDGMENTS

We would like to express our gratitude to Mr. and Ms. Kurihara in Tsukuba for providing the site for the field experiment. We thank Dr. A. Koga (Kyoto University) for his constructive suggestion for this study. We are also grateful to Y. Nakaizumi, D. Nohara, and Dr. W. Mukaimine (University of Tsukuba) for supporting our field experiment. We also thank Drs. D. Kurokawa and T. Miura (The University of Tokyo) for supporting medaka rearing and experiments. We are grateful to National BioResource Project for providing medaka for preliminary experiments. This research was partially supported by JSPS KAKENHI Grant No. 18K06410 to YT.

## LITERATURE CITED

Bowler, Diana E., and Tim G. Benton. 2005. “Causes and Consequences of Animal Dispersal Strategies: Relating Individual Behaviour to Spatial Dynamics.” Biological Reviews of the Cambridge Philosophical Society 80 (2): 205–25.

Coughlan, Neil E., Thomas C. Kelly, John Davenport, and Marcel A.K. Jansen. 2017. “Up, up and Away: Bird-Mediated Ectozoochorous Dispersal between Aquatic Environments.” Freshwater Biology 62 (4): 631–48.

Coughlan, Neil E., Thomas C. Kelly, and Marcel A.K. Jansen. 2017. “‘Step by Step’: High Frequency Short-Distance Epizoochorous Dispersal of Aquatic Macrophytes.” Biological Invasions 19: 625–34.

Faulks, Leanne K., Dean M. Gilligan, and Luciano B. Beheregaray. 2010. “Islands of Water in a Sea of Dry Land: Hydrological Regime Predicts Genetic Diversity and Dispersal in a Widespread Fish from Australias Arid Zone, the Golden Perch (Macquaria Ambigua).” Molecular Ecology 19 (21): 4723–37.

Garcia, Flavien, Ivan Paz-Vinas, Arnaud Gaujard, Julian D. Olden, and Julien Cucherousset. 2023. “Multiple Lines and Levels of Evidence for Avian Zoochory Promoting Fish Colonization of Artificial Lakes.” Biology Letters 19 (3).

Green, Andy J., Ádám Lovas-Kiss, Chevonne Reynolds, Esther Sebastián-González, Giliandro G. Silva, Casper H.A. van Leeuwen, and David M. Wilkinson. 2023. “Dispersal of Aquatic and Terrestrial Organisms by Waterbirds: A Review of Current Knowledge and Future Priorities.” Freshwater Biology 68 (2): 173–90.

Hirsch, Philipp Emanuel, Anouk N’Guyen, Roxane Muller, Irene Adrian-Kalchhauser, and Patricia Burkhardt-Holm. 2018. “Colonizing Islands of Water on Dry Land—on the Passive Dispersal of Fish Eggs by Birds.” Fish and Fisheries 19 (3): 502–10.

Iwamatsu, Takashi. 2004. “Stages of Normal Development in the Medaka Oryzias Latipes.” Mechanisms of Development 121 (7–8): 605–18.

Iwata, Eri, Koutarou Sakamoto, Takuya Ohkouchi, Hideaki Sasaki, Jun Yasuda, and Makito Kobayashi. 2015. “Egg Deposition Sites of Wild Minami-Medaka Oryzias Latipes in an Aquatic Area of a Biotope.” Natural Environmental Science Research 28: 11–21.

Kamide, Sakurako, Yu Kominami, and Makito Kobayashi. 2017. “Preference in Water Depth for Egg Deposition of Female Medaka, Oryzias Latipes.” Natural Environmental Science Research 30: 1–4.

Kobayashi, Makito, Tomohisa Yoritsune, Shohei Suzuki, Ayami Shimizu, Mika Koido, Yutaro Kawaguchi, Youichi Hayakawa, Sayaka Eguchi, Hirofumi Yokota, and Yoshikazu Yamamoto. 2012. “Reproductive Behavior of Wild Medaka in an Outdoor Pond.” Nippon Suisan Gakkaishi (Japanese Edition) 78 (5): 922–33.

Lovas-Kiss, Adam, Orsolya Vincze, Viktor Löki, Felicia Paller-Kapusi, Béla Halasi-Kovács, Gyula Kovács, Andy J. Green, and Balázs András Lukács. 2020. “Experimental Evidence of Dispersal of Invasive Cyprinid Eggs inside Migratory Waterfowl.” Proceedings of the National Academy of Sciences of the United States of America 117 (27): 15397–99.

Nakamura, Takio. 1999. Soratobu Medaka (Flying Medaka in the Sky). Tokyo: POPLAR PUBLISHING CO., LTD.

Senou, Hiroshi. 2013. “Family Adrianichthyidae.” In Fishes of Japan with Pictorial Keys to the Species, edited by Tetsuji Nakabo, 3rd ed., 649–50, 1923–26. Hadano: Tokai University Press.

Silva, Giliandro G., Vinícius Weber, Andy J. Green, Pedro Hoffmann, Vanessa S. Silva, Matheus V. Volcan, Luis Esteban K. Lanés, Cristina Stenert, Martin Reichard, and Leonardo Maltchik. 2019. “Killifish Eggs Can Disperse via Gut Passage through Waterfowl.” Ecology 100 (11): 55–58.

Viana, Duarte S., Luis Santamaría, Thomas C. Michot, and Jordi Figuerola. 2013. “Migratory Strategies of Waterbirds Shape the Continental-Scale Dispersal of Aquatic Organisms.” Ecography 36 (4): 430–38.

