## Appendix S1 for "Freshwater fish egg dispersal by attaching to waterbirds"

**MATERIAL and METHODS**

**Ethical statement**

Experiments in this study followed the animal experiment use guidelines of University of Tsukuba (https://www.md.tsukuba.ac.jp/LabAnimalResCNT/kitei/doubutsukitei.pdf) and that of The University of Tokyo (https://www.u-tokyo.ac.jp/gen01/reiki_int/reiki_honbun/au07404001.html).

**Field experiment**

We constructed two experimental ponds (1 m × 2 m, 1 m distance between ponds) by burying plastic containers (91 cm × 62 cm × 20 cm; three containers in one pond) in a shallow irrigation ditch among paddy fields in Hojo, Tsukuba, Ibaraki, Japan. Akadama sand was laid down on the bottom of the ponds. The water depth of these ponds was about 10 cm. Thirty-six artificial aquatic plants (strap-shaped, about 30 cm, made from polypropylene, Suisaku Co., Ltd., Japan) were placed in the "source pond." Herons and egrets constantly visited paddy fields in the Hojo area for foraging. Since herons and egrets are large waterbirds compared with other birds that came flying in paddy fields, such as spot-billed duck (*Anas zonorhyncha*) and Japanese wagtail (*Motacilla grandis*), and it was thus expected that such large shorebirds would relatively easily carry aquatic plants, we focused on them. It is known that herons and egrets eat fish, frogs, crustaceans, insects, and other small animals (Ogasawara, Abe, and Naito 1982; Tojo 1996). Therefore, to attract herons and egrets, small baitfish (topmouth gudgeon, *Pseudorasbora parva*, continental rosy bittering, *Rhodeus ocellatus ocellatus*, and japanese weatherfish, *Misgurnus sp.*, collected in Tsukuba city) were added to these ponds, and replaced in response to their absence (about 10 individuals/plastic container). Arrivals of animals and their behaviors around ponds were recorded by two motion capture camera traps (CMS-SC03GY, SANWA SUPPLY Inc., Okayama, Japan). The recording parameter of the cameras was to record 60 seconds per each bird/animal arrival (detected by infrared sensors) for the whole day during the experimental period. Also, Supplemental video 3 was recorded using a digital camera (PowerShot SX530HS, Canon, Tokyo, Japan) on May 23, 2019, in Hojo, Tsukuba, Ibaraki, Japan.

**Egg dehydration experiment**

Parental medaka were collected using a hand-net at the pond in the Yata River system, Tsukuba, Ibaraki, Japan in October 2023. Each Pair of medaka was reared for collecting eggs in a separate tank (32 cm× 19 cm× 24 cm) with filtering and aeration at a water temperature of 26°C; light period: dark period (L: D) =14h:10h. We also used individuals provided from National BioResource Project Medaka (strain ID: WS226, WS227) for preliminary experiments. Egg clutches were collected every morning and stored in freshwater supplemented with 0.1% methylene blue (rearing water) at 25°C, L: D=12h:12h in a six-well dish in an incubator (MTI-201, Tokyo Rikakikai Co., Ltd.,

Japan). Four eggs from each clutch were kept in rearing water as a control. If the control eggs did not hatch, we removed the data based on this clutch. Since control eggs did not hatch or rearing water dried up accidentally, three data points were removed from the dataset. Clutches one day after spawning were used for the experiment. Four eggs were attached to a 5-cm piece of Anacharis *Egeria densa*. Then, eggs were exposed to the air with 75% relative humidity and 25°C in an incubator (MIR-154, SANYO Electric Co. Ltd., Japan). Humidity was maintained using a saturated saline solution in the incubator. Temperature and humidity were checked using Thermo-hygrometer (IBS-TH1-mini, Inkbird, China). Such environmental conditions were similar to the average humidity and temperature recorded in Tsukuba by the Japan Meteorological Agency in July, which was medaka's reproductive season (Iwamatsu 2004). Seven exposure times were applied (0, 6, 12, 15, 18, 21, and 24 hours). After treatment, eggs were kept in rearing water until they hatch. Usually, medaka eggs hatch for about 10 days at 26°C (Iwamatsu 2004). Each treatment was performed from seven to 12 biological replicates. In addition, four eggs were exposed to the air with (attached to 5 cm of Anacharis) and without (placed on the sliding glass) aquatic plants to evaluate the importance of moisture retention by attachment to aquatic plants for egg survival (N = 8 biological replicates).

**Data analysis**

Statistical analysis was performed using R 4.3.1 (R Core Team 2023). We examined the association of hatching rate and the period in the air using probit regression model following the previous study (Banha and Anasta 2012). Median lethal time (the time of 50% mortality or $LD_{50}$) of exposed air was calculated based on hatching rate. Also, hatching rates with and without aquatic plants were compared using Wilcoxon signed-rank test.

**Supplemental References**

Banha, Filipe, and Pedro Manuel Anasta. 2012. "Waterbird-Mediated Passive Dispersal of River Shrimp *Athyaephyra Desmaresti*." *Hydrobiologia* 694: 197–204.

Iwamatsu, Takashi. 2004. "Stages of Normal Development in the Medaka Oryzias Latipes." *Mechanisms of Development* 121 (7–8): 605–18.

Ogasawara, Ko, Kaname Abe, and Toshihiko Naito. 1982. "Ecological Study of Grey Heron in Oga Peninsula, Akita Prefecture." *Journal of the Yamashina Institute for Ornithology* 14 (2–3): 232–45.

R Core Team. 2023. "R: A Language and Environment for Statistical Computing. R Foundation for Statistical Computing, Vienna, Austria."

Tojo, Hitoshi. 1996. "Habitat Selection, Foraging Behaviour and Prey of Five Heron Species in Japan." *Japanese Journal of Ornithology* 45 (3): 141–58.

**Captions of supplemental videos**

**Video S1.**

A grey heron hooked artificial aquatic plants on its leg and was walking in the sink pond (experimental pond). This video was taken by a motion capture camera trap. Recorded on Dec. 5, 2019, in Hojo, Tsukuba, Ibaraki, Japan.

Videographer: Akifumi Yao

**Video S2.**

A grey heron hooked artificial aquatic plants on its leg and flew away from the sink pond (experimental pond). This video was taken by a motion capture camera trap. Recorded on Dec. 5, 2019, in Hojo, Tsukuba, Ibaraki, Japan.

Videographer: Akifumi Yao

**Video S3.**

A grey heron walked between paddy fields tangled clump of algae on its leg. Recorded on May 23, 2019, in Hojo, Tsukuba, Ibaraki, Japan.

Videographer: Miyuki Mashiko
